# Rapid extraction and kinetic analysis of protein complexes from single cells

**DOI:** 10.1101/2021.07.07.451367

**Authors:** Sena Sarıkaya, Daniel J. Dickinson

## Abstract

Proteins contribute to cell biology by forming dynamic, regulated interactions, and measuring these interactions is a foundational approach in biochemistry. We present a rapid, quantitative *in vivo* assay for protein-protein interactions, based on optical cell lysis followed by time-resolved single-molecule analysis of protein complex binding to an antibody-coated substrate. We show that our approach has better reproducibility, higher dynamic range, and lower background than previous single-molecule pull-down assays. Furthermore, we demonstrate that by monitoring cellular protein complexes over time after cell lysis, we can measure the dissociation rate constant of a cellular protein complex, providing information about binding affinity and kinetics. Our dynamic single-cell, single-molecule pull-down method thus approaches the biochemical precision that is often sought from *in vitro* assays, while being applicable to native protein complexes isolated from single cells *in vivo*.

## Introduction

Protein-protein interactions are fundamental determinants of protein function. By interacting and forming complexes, proteins assemble macromolecular machines that carry out a myriad of activities within cells. A major goal of experimental biology is to observe, measure, and characterize these interactions, and to understand how they are controlled and contribute to cell behavior.

For protein-protein interaction assays to be maximally informative, they should ideally be performed *in vivo –* that is, using natively expressed proteins engaged in their normal cellular functions. Unfortunately, traditional assays such as co-immunoprecipitation usually rely on protein overexpression and do not typically yield quantitative information about protein complex abundance or binding affinity. Therefore, investigators interested measuring quantitative binding parameters such as the dissociation constant (K_d_) or binding / unbinding rate constants (k_on_ and k_off_) generally turn to *in vitro* approaches with purified proteins (1, 2). While such approaches provide an unparalleled degree of experimental control and can yield quantitative descriptions of protein-protein interactions, they may not always recapitulate the physiological state of the proteins under study.

In the past decade, technical advancements from our group (3) and others (4–7) have begun to improve the potential of cell-based assays to yield quantitative information about protein complexes. A pioneering study from Jain and colleagues (4) showed that protein-protein interactions could be assayed in cell lysates by immunoprecipitating fluorescently tagged proteins onto microscope coverslips and imaging the resulting immunocomplexes using single-molecule TIRF microscopy, an approach termed <u>si</u>ngle-<u>m</u>olecule <u>pull</u>-down (SiMPull). Because SiMPull counts individual protein complexes, it can provide a digital, quantitative measurement of the fraction of proteins in complex in a given sample. Subsequent work from the Yoon lab (6, 7) further developed this approach by pulling down a tagged protein and then adding a second cell lysate carrying a tagged binding partner; by monitoring these reactions over time, they were able to extract kinetic parameters describing the binding. However, the protein complexes studied in this approach were formed on the coverslip surface, and thus did not represent native cellular interactions (6).

Building on this earlier work, we developed a method termed single-cell, single-molecule pull-down (sc-SiMPull) (3) for studying near-*in vivo* protein interactions from single-cell embryos. To develop this approach, we made three key changes to previous SiMPull approaches. First, we miniaturized SiMPull by lysing single cells inside nanoliter-volume microfluidic channels. Microfluidic lysis reduced the amount of input material required, enabling single-cell analysis. The microfluidic approach also allowed faster sample processing; samples were analyzed minutes instead of hours after lysis. Second, we developed and used genome editing technology (8, 9) to insert fluorescent tags directly into the native genes encoding the proteins under study, eliminating the need for overexpression and ensuring that any protein complexes we detected were formed by endogenous proteins. Third, we developed freely available, user-friendly image analysis software that eliminates the need for manual counting of molecular complexes and enables a user to obtain data from hundreds to thousands of molecules per cell (3). Together, these improvements enabled us to detect changes in native protein complex abundance that occurred over the span of a few minutes during development of the *Caenorhabditis elegans* zygote (3). We showed that the PAR-3 protein, which is critical for embryonic cell polarization, is found in oligomers that dynamically assemble and disassemble during the first cell cycle. PAR-3 oligomerization cooperatively promotes its association with the key kinase aPKC, and thereby enables aPKC transport to the anterior side of the cell to establish the anterior-posterior axis (3). Despite enabling these important molecular insights, our initial sc-SiMPull approach still had significant limitations. It relied on a manual mechanical method to lyse cells, hindering reproducibility and precise embryo staging. In addition, the time from cell lysis to data collection was still several minutes, which obscures kinetic information and precludes detecting weak or transient complexes that may dissociate within seconds after cell lysis.

Here, we further improve upon the sc-SiMPull approach by incorporating near-instantaneous, on-chip cell lysis by a pulsed infrared laser. We demonstrate that laser lysis is compatible with sc-SiMPull; it does not photobleach the fluorescent labels that are required for single-molecule imaging, nor does it detectably damage cellular protein complexes. We further show that, by imaging captured protein complexes over time and monitoring their dissociation, we can measure the kinetics of protein complex dissociation and obtain important information about the stability of protein-protein interactions.

## Results

### Incorporating laser-induced cell lysis into an sc-SiMPull assay

We use *C. elegans* zygotes as model cells throughout this work because they are easy to isolate and image, develop via a stereotyped series of morphologically distinct stages (10) and can be easily manipulated via CRISPR to carry fluorescent tags on endogenous proteins (9, 11, 12). In the standard sc-SiMPull assays we previously described (3), an individual zygote is first selected and trapped in a funnel-shaped microfluidic device made of polydimethylsiloxane (PDMS) (Figure 1A, left). The cell is staged via manual observation on a stereomicroscope and then lysed mechanically, by pushing down on the PDMS surface (Figure 1A, top). Upon lysis, cellular proteins diffuse within the microfluidic channel and bind to antibodies, which are immobilized on a coverslip that forms the floor of the device (Figure 1A, right center). After cell lysis the device is transferred to a TIRF microscope and single molecule images are acquired. Transferring the device to the TIRF microscope is the rate-limiting step in this assay; moving the device, focusing, configuring and starting data acquisition takes a minimum of 2-3 minutes. During this time, protein complexes can rearrange, so that the results represent the pre-lysis state only in the case of stable protein complexes. Moreover, the manual nature of the staging and mechanical lysis introduces unwanted variability between experiments and experimenters.

**Figure 1:**
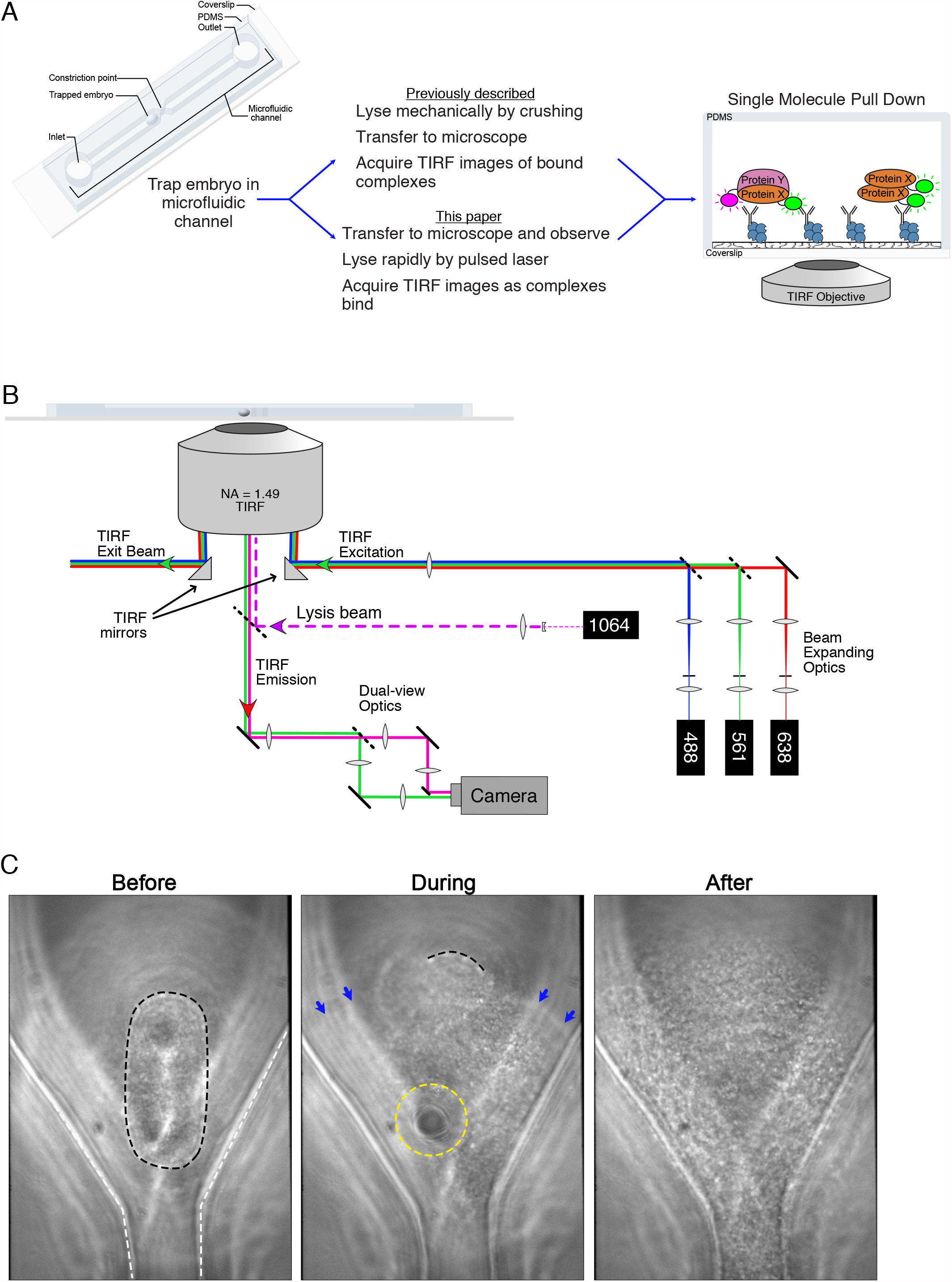
Laser-induced lysis for single-cell, single-molecule pull-down. A) Overview of sc-SiMPull using mechanical vs. laser-induced cell lysis. In both cases, the workflow begins by trapping an embryo in a PDMS microfluidic device with a volume of a few nL (left). After cell lysis, proteins are captured by antibodies immobilized on the coverslip surface and are detected using single-molecule TIRF imaging (right). In the conventional workflow (center top), the cell is lysed before being transferred to the TIRF microscope (3); our new workflow (center bottom) uses a laser to perform lysis directly at the imaging microscope, allowing more rapid and reproducible data collection (see below). B) Schematic of the microscope used to perform these experiments. TIRF illumination is performed using micromirrors located below the objective, allowing multi-wavelength, dichroic-free single-molecule imaging (17, 18). Fluorescence emission is detected with a dual-view emission path design for simultaneous multicolor imaging. We added a pulsed 1064 nm laser to this setup, focused at the sample plane, to induce cell lysis via formation of a localized plasma. C) Brightfield images of a *C. elegans* zygote before, during and after laser-induced lysis. The black dotted line outlines the cell membrane of the single-cell embryo. The white dotted line outlines the edges of the microfluidic channel. The yellow dotted line outlines the edges of the cavitation bubble. Blue arrows point to the edges of the shock wave as it moves through the channel. See also Movie S1.

To overcome these limitations, we turned to cell lysis induced by a passively Q-switched Nd:YAG pulsed laser. These lasers are compact, relatively inexpensive, and produce pulses of laser light lasting several hundred picoseconds. When focused to a diffraction-limited spot by a high numerical aperture objective lens, these lasers produce a localized plasma, which in turn generates a shock wave and a cavitation bubble in aqueous medium (13–15). Lysis occurs within a few microseconds after irradiation and is thought to occur as a result of mechanical disruption of the cell membrane by the shock wave and cavitation bubble, and not by the laser light *per se* (13–15). Since formation of a shock wave and cavitation bubble is relatively independent of laser wavelength (16), we chose to use an infrared 1064 nm laser for cell lysis to avoid interference with subsequent fluorescence-based assays.

We installed and configured the lysis laser on a multi-wavelength TIRF microscope based on a micromirror geometry (17, 18). In a micromirror TIRF instrument, a small mirror (“micromirror”) is positioned directly below the back aperture of the objective lens to deliver TIRF excitation light to the sample, and a second micromirror collects the reflected TIRF beam and delivers it to a beam dump (Figure 1B). The fluorescence emission passes between the two micromirrors to a multiwavelength imaging system (dual-view optics in Figure 1B). Micromirror TIRF enables straightforward single-molecule imaging using multiple wavelengths simultaneously, which in turn enables kinetic measurements of protein complex formation and dissociation (17, 18). We added a short-pass dichroic mirror below the micromirrors to reflect the infrared lysis beam towards the sample without interfering with fluorescence detection in visible wavelengths (Figure 1B). The lysis beam was expanded and collimated to fill the back aperture of the objective, thereby focusing to a diffraction-limited spot in the sample plane. A single shot from the laser produced a shock wave and cavitation bubble, which was sufficient to lyse a *C. elegans* zygote (Figure 1C and Movie S1).

### Laser lysis is compatible with SiMPull and improves reproducibility

We performed control experiments to determine whether laser lysis was compatible with single-molecule fluorescence assays such as sc-SiMPull. First, we checked whether the lysis laser induced photobleaching of mNeonGreen (mNG) or JaneliaFluor 646 (JF_646_), which are the two fluorophores we use most commonly in sc-SiMPull assays. To check for photobleaching, we lysed zygotes expressing an mNG::HaloTag fusion protein that was labeled with Halo-JF_646_, captured the fusion protein with anti-mNG nanobodies and analyzed the colocalization between green (mNG) and far-red (JF_646_) signals (Figure 2A, top). Since mNG and HaloTag are covalently attached to one another in this experiment, 100% of the green and far-red signals should colocalize in principle. In practice, as we demonstrated in our previous experiments, less than 100% colocalization is observed due to incomplete maturation of mNG fluorescent protein and incomplete labeling of the HaloTag with its ligand dye (3). The actual extent of colocalization is a measurement of the efficiency with which each fluorescent tag is detected. Photobleaching would appear as a reduced detection efficiency for the bleached dye. Using our previous method of lysing zyogtes mechanically by pressing on the microfluidic device with the tip of a pencil, we detected 63% ± 8% (mean ± 95%CI) of mNG molecules and 72% ± 4% of JF_646_ molecules (Figure 2A, bottom), consistent with our previous results (3). We detected 59% ± 4% of mNG molecules and 73% ± 3% of JF_646_ molecules when we lysed cells with the laser, which is indistinguishable from the results of mechanical lysis (Figure 2A). We conclude that laser lysis does not detectably photobleach mNG or JF_646_ fluorescent labels (3).

**Figure 2:**
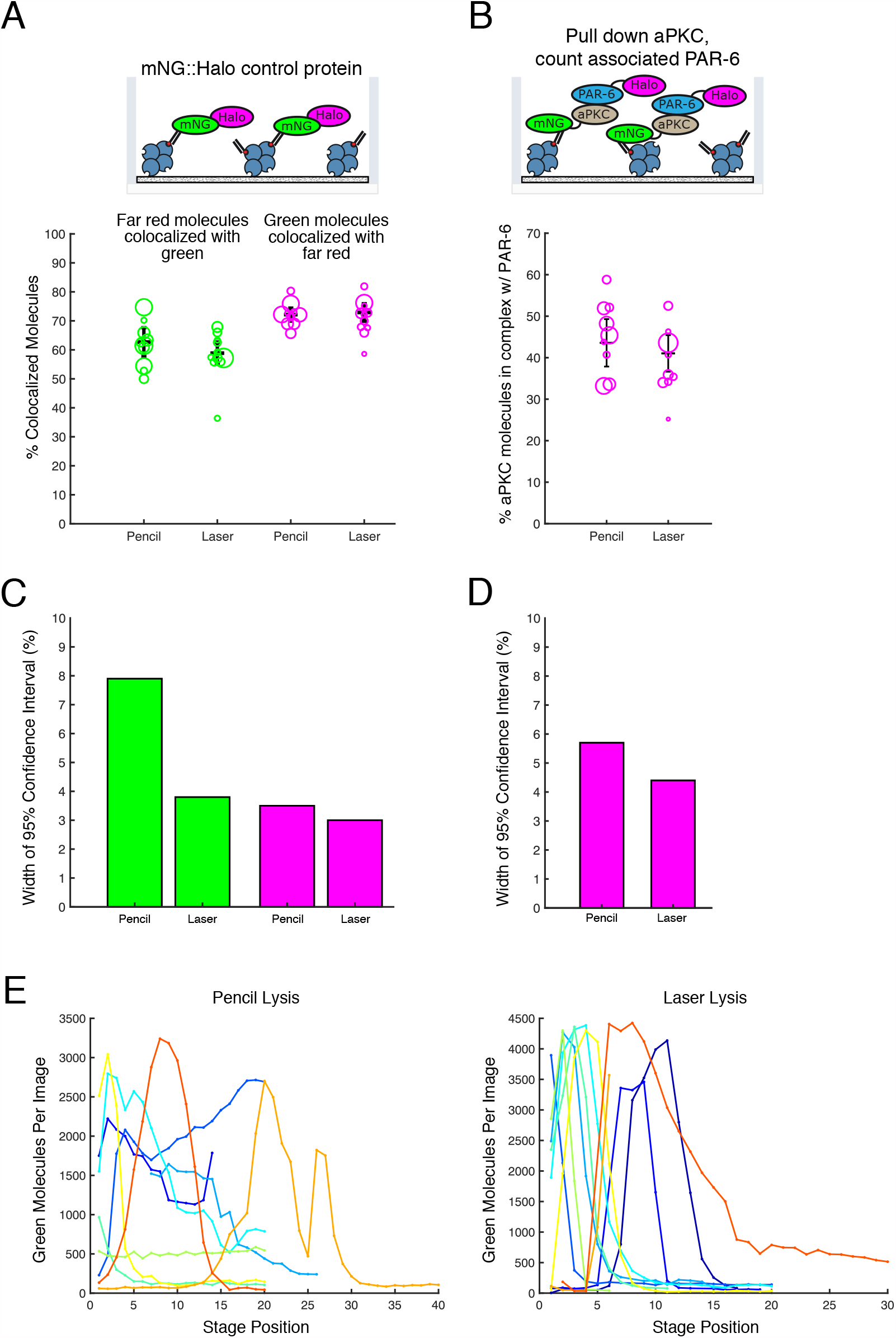
Laser lysis is compatible with SiMPull and improves reproducibility. A) Comparison of laser-induced and mechanical (“pencil”) lysis using an mNG::HaloTag control protein, where the two tags are covalently fused as illustrated at the top. In the chart at the bottom, each circle represents a measurement from a single cell, and the size of each circle represents the number of molecules counted in that experiment. Regardless of the lysis method, we detect ∼60% of mNG molecules and ∼75% of HaloTag molecules in a typical positive control experiment. N_pencil_=10; N_laser_=11. B) Comparison of laser-induced and mechanical lysis for analysis of the aPKC/PAR-6 interaction (top). Each circle represents a measurement from a single cell, and the size of each circle represents the number of molecules counted in that experiment (bottom). N_pencil_=9; N_laser_=9. C,D) Widths of the 95% confidence intervals of the weighted means for each dataset presented in (A) and (B). E) Distribution of mNG::HaloTag molecules along the length of the microfluidic channel, measured by acquiring images at several stage positions. Each curve shows the results from a single experiment. Stage positions are 150 µm apart.

We next asked whether protein-protein interactions are preserved after laser lysis. Laser lysis has been proposed to be non-damaging to cellular proteins (19, 20). Indeed, protein damage is not expected in our setup because the laser focus (where plasma formation occurs) is located at the glass–water interface, which is outside the cell, and lysis occurs due to the mechanical effects of cavitation bubble formation. Nevertheless, we wanted to directly test whether a known protein complex was stable through the laser lysis process. We therefore examined the interaction between aPKC and PAR-6, two proteins that are known to form a complex in *C. elegans* zygotes (3, 21, 22). We lysed zygotes from a strain carrying mNG::aPKC and PAR-6::Halo tagged at their endogenous genomic loci (3), captured mNG::aPKC with an anti-mNG nanobody, and measured the fraction of mNG::aPKC molecules associated with PAR-6::Halo (Figure 2B, top). We observed an equivalent fraction of molecules in complex regardless of the lysis method (Figure 2B, bottom) and the results again agree with our previous report (3). We conclude that laser lysis does not disrupt the aPKC/PAR-6 interaction. Although these data do not rule out that some protein complexes might be affected by laser lysis, they argue against a large-scale disruption of protein-protein interactions by our laser setup (19, 20).

Interestingly, when analyzing these control experiments, we observed a small but consistent trend towards lower cell-to-cell variability in our SiMPull data when cells were lysed by the pulsed laser (Figure 2C–D). We also found that, when we analyzed the distribution of captured molecules along the length of the microfluidic channel, cells lysed by the laser produced a narrower and more consistent peak of signal near the site of lysis than cells lysed manually (Figure 2E). Together, these observations suggest that laser lysis improves the reproducibility of sc-SiMPull experiments. We also note that when performing laser lysis experiments, we routinely acquire an image of the cell immediately before lysis, which will lend additional confidence to experiments that require precise staging (e.g. (3)). We conclude that laser lysis improves the speed, precision and reproducibility of sc-SiMPull experiments without otherwise affecting the results.

### Dynamic analysis of cellular protein complexes after single-cell lysis

We next considered whether rapid lysis might enable us to gain kinetic information that would not be available using other methods. In the typical workflow that we have used up to this point, we lyse a cell, incubate the lysate for several minutes, and then analyze molecules that are bound to the coverslip surface. We refer to this approach as a “static” workflow (Figure 3A, left). Static sc-SiMPull experiments can provide information about the abundance and stoichiometry of protein complexes that are stable enough to survive several minutes of incubation after cell lysis, but static experiments do not provide any kinetic information and are not applicable to transient complexes. We envisioned an alternative “dynamic” workflow in which we would begin acquiring data immediately (within seconds) after lysis and observe the progression of the binding reaction between immobilized antibodies and a protein complex of interest (Figure 3A, right). We show below how dynamic experiments provide kinetic information about the stability and dynamics of protein complexes retrieved directly from cells.

**Figure 3:**
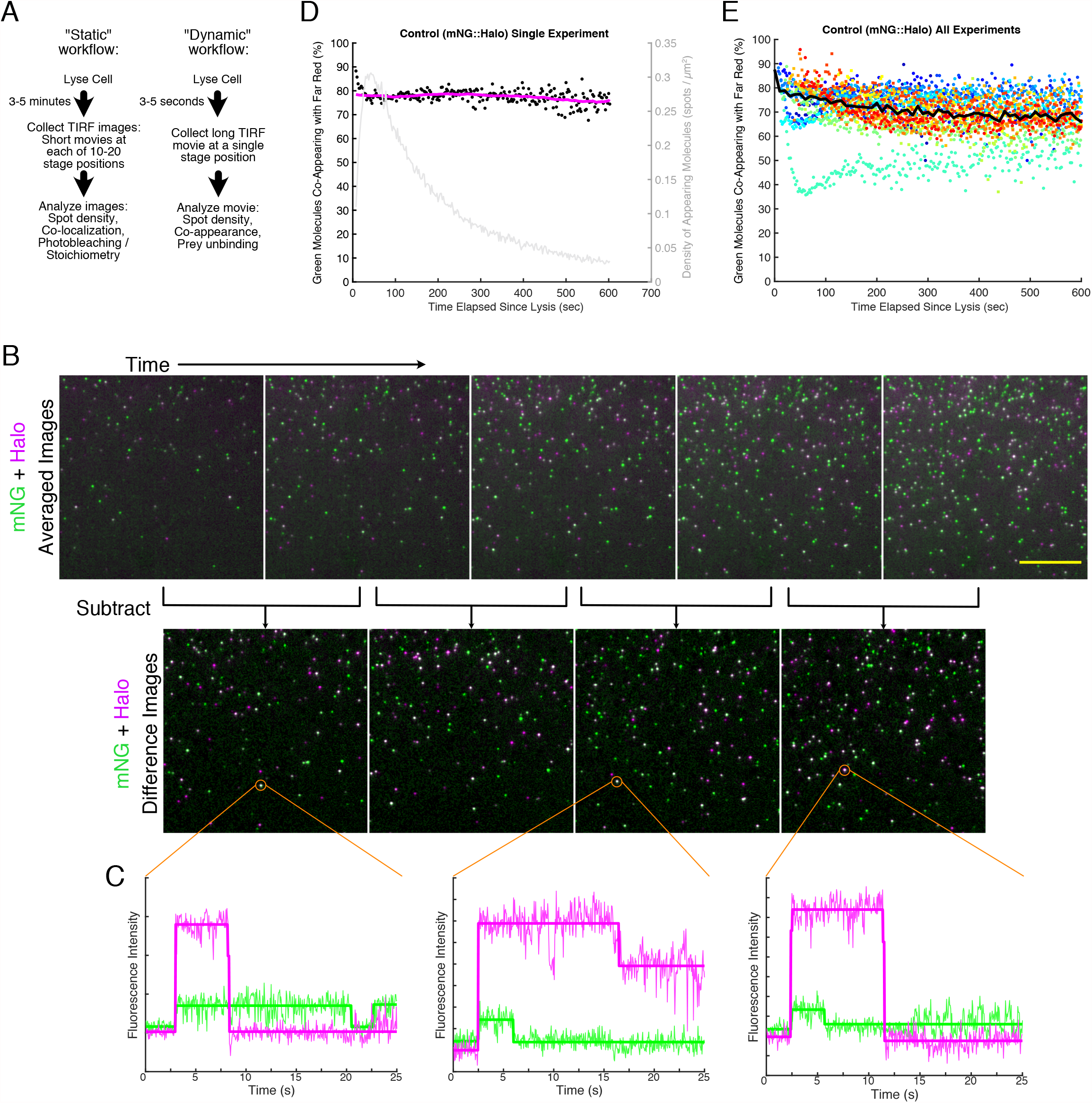
A dynamic sc-SiMPull workflow. A) Comparison of our previous “static” workflow (left) and the new “dynamic” workflow (right) enabled by laser lysis. B) Illustration of a typical dynamic dataset and our data processing approach. Top row: average intensity projections of consecutive 50-frame windows near the beginning of a dynamic dataset. Molecules appear at the coverslip over time following cell lysis. Each image corresponds to 2.5s worth of data. Scale bar represents 10 µm. Second row: difference images obtained by subtracting each average intensity projection from the one that follows. Difference images are segmented to identify appearing molecules. C) Example intensity vs. time traces for co-appearing signals. The three indicated molecules were all classified as co-appearing based on the simultaneous increase in signal in both channels around 3s after the start of the trace. D) Results of one dynamic sc-SiMPull experiment with the mNG::HaloTag control protein. Black points show individual measurements of the fraction of molecules that co-appeared; each point corresponds to one 50-frame (2.5s) window of the data. The magenta curve is a moving average of these data. The gray curve shows the density of appearing spots as a function of time, plotted on the right Y axis. E) Results of a complete set of experiments with the mNG::HaloTag control protein. Each color represents individual measurements for one sample, and the black curve is a moving average across all the data. Circles represent data that were acquired at 50 mW laser power, and squares represent data that were acquired at 5 mW laser power. The sea-green dataset shows anomalously low co-appearance but was included here for completeness. N=14.

To realize this approach, we first studied the control mNG::HaloTag fusion protein. We lysed embryos carrying this protein, immediately moved to an adjacent stage position, and observed the coverslip surface over time using TIRF. Acquisition typically started 5-10 seconds after cell lysis. In the resulting movies, fluorescent molecules began to appear over time in a region of the device that initially had been empty (Figure 3B, top row and Movie S2). As expected, many of these molecules were fluorescent in both the green (mNG) and far-red (HaloTag) channels. We refer to the simultaneous appearance of a fluorescent signal in both channels as “co-appearance.” Co-appearance is the dynamic analog of the colocalization between labels that we measure in a static experiment. When two fluorescently tagged molecules co-appear, we infer that they are in a complex. Co-appearance is a more stringent metric for protein-protein interaction than colocalization because it requires that signals occur together not only in the 2D space of the image, but also in time.

We developed an open-source MATLAB package to extract, quantify and visualize co-appearance data from time-lapse TIRF movies of single-molecule appearances (see Methods for details). In brief, our analysis pipeline consists of two steps: Molecule detection and co-appearance analysis. First, for molecule detection, each movie is divided into windows of 20-50 frames, and images in each window are averaged to increase signal:noise (Figure 3B, top row). To identify newly appearing spots, pairs of adjacent averaged images are subtracted, producing difference images such that visible signals are spots that appeared in that window (Figure 3B, bottom row). The difference images are segmented to identify the locations of newly appearing spots. Second, for co-appearance analysis, an intensity vs. time trace is extracted for each spot (from the raw, not averaged, data) (Figure 3C). The software processes each trace using a Bayesian framework to detect changes in signal intensity (23) and calculates a mean intensity for each segment of each trace (Figure 3C, bold lines). Traces in which both channels show a simultaneous increase in intensity (as in Figure 3C) are considered co-appearing. A gallery of traces, showing examples of different behaviors, is presented in Figure S1. While developing this analysis platform, we identified missed or erroneous co-appearance events caused by photoblinking (Figure S2A) and by molecule densities that were too low (Figure S2B) or too high (Figure S2C), and our software incorporates optional settings to filter out these artifacts automatically.

Since co-appearance is a new metric for protein-protein interactions, we benchmarked our analysis pipeline using the control mNG::HaloTag protein. We observed that 70-80% of mNG molecules co-appeared with HaloTag (Figure 3D-E), in agreement with our static data using the same construct and fluorophores (Figure 2B), confirming that our co-appearance analysis successfully detected proteins that are associated with one another. We also found two other notable features of the co-appearance metric. First, co-appearance was less sensitive to molecular density than static colocalization: We could accurately measure co-appearance at densities up to at least 0.8 molecules present per µm^2^ (Figure S2C-D), while colocalization measurements are reliable only up to densities of ∼0.3 molecules per µm^2^ (3). Our ability to detect co-appearance did suffer at extremely high densities (>0.8 molecules per µm^2^), likely due the difficulty of extracting accurate intensity vs. time traces from crowded images (Figure S2C-D). Second, co-appearance measurements had lower background: a pair of proteins that were not expected to interact co-appeared less than 2% of the time (Figure S2E), whereas static datasets can exhibit non-specific colocalization of 5-10% of spots depending on the density of molecules on the coverslip (3, 31). We conclude that co-appearance of molecules in a single molecule pull-down assay is a robust, accurate and specific metric of protein-protein interactions.

### Dynamic analysis provides information about protein-protein interaction kinetics

We predicted that dynamic, time-resolved measurements of protein-protein interactions in fresh cell lysates could provide a wealth of information about the kinetics and strength of these interactions. To explore this idea, we studied the aPKC/PAR-6 complex. When we lysed a zygote and captured mNG::aPKC molecules with an anti-mNG nanobody, we observed co-appearing PAR-6::HaloTag molecules, indicating that aPKC and PAR-6 are in complex, as expected (Movie S3). The initial fraction of co-appearing signals was 40–50%, consistent with our results from static data (compare Figure 4B to Figure 2B). Interestingly, however, we found that the fraction of co-appearing spots declined over time (Figure 4A–B). This decline was not due to photobleaching, because control mNG::HaloTag samples imaged on the same days and under identical conditions did not show a similar decline in co-appearance over time (compare Figure 4B to Figure 4C). We therefore deduce that the decline in co-appearance is due to dissociation of the aPKC/PAR-6 complex in solution over time following cell lysis.

**Figure 4:**
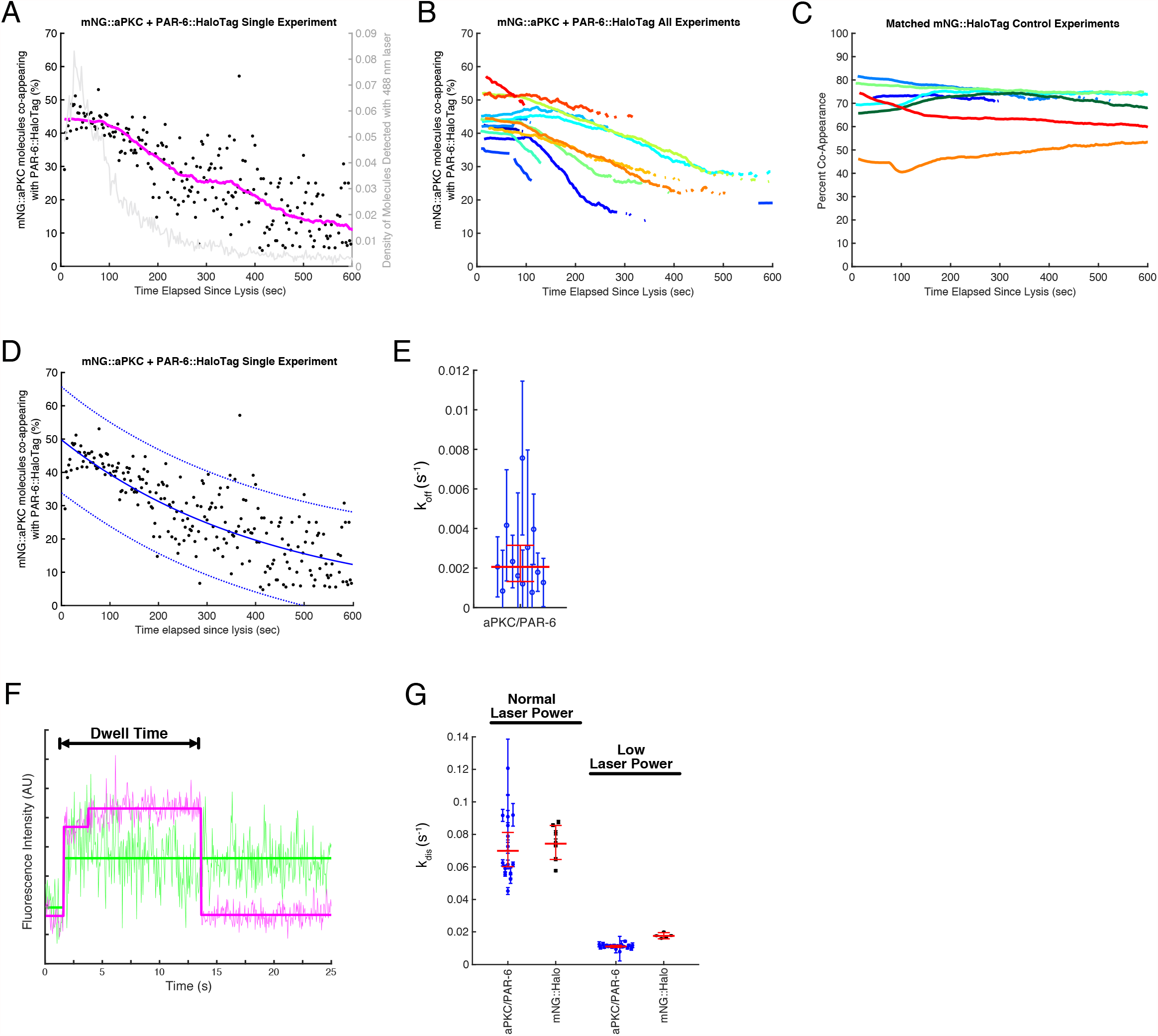
Kinetic analysis of the aPKC/PAR-6 complex released from cells. A) Results of one dynamic sc-SiMPull experiment with mNG::aPKC as the bait protein and PAR-6::HaloTag as the prey protein. Black points show individual measurements of the fraction of molecules that co-appeared; each point corresponds to one 50-frame (2.5s) window of the data. The magenta curve is a moving average of these data. The gray curve shows the density of appearing spots as a function of time, plotted on the right Y axis. B) Co-appearance vs. time for the full set of aPKC/PAR-6 experiments. For clarity, only the trendlines (equivalent to the magenta trace in (A)) are shown. N=13. C) Co-appearance vs. time for a matched set of mNG::HaloTag control experiments. These samples were collected and imaged on the same days and with identical laser power to the aPKC/PAR-6 samples in (B). N=7. D) The same data as in (A) fit to an exponential decay function (blue solid curve). Blue dashed curves show the 95% confidence intervals of the fit. E) Estimates of the dissociation rate constant *k*_*off*_ from exponential fits to each individual datasets (as in (D)). Each point shows the result of one experiment with its 95% confidence interval. The horizontal red line and error bars represent the geometric mean across all samples, with its 95% confidence interval. N=12. F) Intensity vs. time trace for a representative mNG::aPKC molecule that co-appeared with PAR-6::HaloTag, illustrating the dwell time that was measured to determine the disappearance rate constant *k*_*disappear*_ from dwell time distribution analysis. G) Measurements of *k*_*disappear*_ for aPKC/PAR-6 samples and control mNG::HaloTag samples at two different laser powers. Each point shows the results of one experiment, with the error bars indicating the 95% credible interval from the Bayesian dwell time distribution analysis (28). The horizontal red line and error bars represent the geometric mean across all samples, with its 95% confidence interval. N_aPKC/PAR-6_=15; N_mNG::Halo_=7; N_aPKC/PAR-6 Low Laser_=19; N_mNG::Halo Low Laser_=5.

To quantify the stability of the aPKC/PAR-6 complex, we fit the co-appearance data from each sample to an exponential decay (Figure 4D shows an example fit) to extract the apparent decay rate constant *k*_*decay*_. We derived a relationship between *k*_*decay*_ and the dissociation rate constant k_off_ (see methods) and used this relationship to estimate *k*_*off*_ for each sample. We obtained an estimated *k*_*off*_ = 2.0×10^−3^ s^-1^ (geometric mean; 95% confidence interval from 1.3×10^−3^ s^-1^ to 3.2×10^−3^ s^-1^) from these measurements (Figure 4E). The equilibrium dissociation constant *K*_*D*_ is related to *k*_*off*_ by the equation 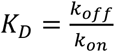. Since *k*_*on*_ values for protein-protein association reactions are typically 10^6^ – 10^7^ M^-1^ s^-1^ (1), our results imply that the aPKC/PAR-6 interaction most likely has a *K*_*D*_ in the range of 0.2–2 nM. This affinity indicates very tight binding, comparable to typical high affinity antibody-antigen interactions. This result is consistent with the known robust association between aPKC and PAR-6 (3, 21, 22, 24–27).

An alternative way to estimate *k*_*off*_ for a protein complex is to measure the lifetime of the bound state, taking advantage of the ability to directly observe binding and unbinding events in single-molecule TIRF experiments (17, 18, 28). Our automated analysis allowed us to readily measure dwell times – defined as the time from complex co-appearance to prey protein disappearance (Figure 4F) – from thousands of aPKC/PAR-6 complexes. From the distribution of dwell time measurements, it is straightforward to calculate a disappearance rate constant *k*_*disappear*_ and its confidence interval (28). This observed rate constant will be faster than the true *k*_*off*_ because prey molecules can disappear due to photobleaching in addition to dissociation of the complex. The disappearance rate constant is the sum of the rate constants for complex dissociation and photobleaching: *k*_*disappear*_ *= k*_*off*_ *+ k*_*bleach*_. Fortunately, we can measure *k*_*bleach*_ from parallel experiments with the covalent mNG::HaloTag fusion protein, for which *k*_*off*_ *=* 0. We performed this analysis and found *k*_*disappear*_ = 7.0×10^−2^ s^-1^ (95%CI from 6.0×10^−2^ s^-1^ to 8.1×10^−2^ s^-1^) for the aPKC/PAR-6 complex and *k*_*bleach*_ = 7.4×10^−2^ s^-1^ (95%CI from 6.5×10^−2^ s^-1^ to 8.5×10^−2^ s^-1^) for mNG::HaloTag samples imaged in parallel (Figure 4G). *k*_*bleach*_ and *k*_*disappear*_ are equal within experimental error, indicating that the dwell times of aPKC/PAR-6 complexes are dominated by photobleaching rather than complex dissociation. This finding is consistent with the fact that *k*_*bleach*_ is more than 30-fold faster than the *k*_*off*_ measured from exponential curve fitting, above; under such conditions, a large majority of prey molecules will photobleach before they dissociate. Next, we attempted to reduce *k*_*bleach*_ by lowering the laser power used to image the PAR-6::HaloTag prey protein. Reducing the laser power strongly reduced the rate of molecule disappearance (Figure 4G), confirming that the disappearance of PAR-6::HaloTag in these experiments is dominated by photobleaching. Under lower laser power conditions, we found *k*_*disappear*_ = 1.11×10^−2^ s^-1^ (95%CI from 1.05×10^−2^ s^-1^ to 1.18×10^−2^ s^-1^), which is reduced compared to our normal imaging conditions but still an order of magnitude greater than our estimate of *k*_*off*_ from curve fitting, above (Figures 4D–E). Even under low laser power conditions, *k*_*disappear*_ and *k*_*bleach*_ were similar; we could not detect a faster *k*_*disappear*_ (which would indicate unbinding) for the aPKC/PAR-6 complex compared to the control protein (Figure 4G). This result validates our conclusion that the aPKC/PAR-6 complex dissociates very slowly due to high-affinity binding. Although the aPKC/PAR-6 complex has a *k*_*off*_ that is too slow to be estimated from our dwell time measurements, the results still show that *k*_*disappear*_ can be readily measured from our data. We anticipate that dwell time distribution analysis will be useful for more transient complexes, which will undergo measurable dissociation on a timescale comparable to (or faster than) photobleaching. Conversely, we expect that it will be more difficult to measure *k*_*off*_ from curve fitting for transient interactions, because such interactions might not generate long enough co-appearance vs. time traces to enable robust fitting. Therefore, the curve fitting and dwell time approaches to measure *k*_*off*_ are complementary: the curve fitting approach is best suited to stable interactions such as aPKC/PAR-6, while the dwell time approach will be best for more transient interactions.

## Discussion

Regulated assembly of protein complexes is a major mechanism by which cells process information for signal transduction and control of cell behavior. To study the regulation of protein-protein interactions, there is a need for new tools that can measure these interactions rapidly, quantitatively, and *in vivo*. Here, we have addressed this need by developing instrumentation and software for rapid single-cell, single-molecule pull-down assays. Our dynamic sc-SiMPull approach allows us to measure the strength of an interaction (in the form of a *k*_*off*_) using native protein complexes isolated directly from living cells. By using CRISPR to label endogenous proteins, we work exclusively with proteins at their native expression levels. Although our assays do not yet reach the degree of precision and control that is possible *in vitro* with purified proteins, our approach has the strength that it is applicable to proteins and protein complexes that are post-translationally modified, difficult to purify, or both.

We used dynamic sc-SiMPull to study the dynamics of the aPKC/PAR-6 complex from *C. elegans* zygotes and measured a *k*_*off*_ in the range of 10^−3^ s^-1^, suggesting low-nanomolar binding affinity. The tight binding affinity we have estimated is consistent with the propensity of PAR-6 to aggregate when not bound to aPKC (29), and with the observation that *in vivo*, PAR-6 is degraded when aPKC is depleted by RNAi (21). To our knowledge, this is the first direct estimate of the affinity of this interaction, likely because PAR-6 is challenging to purify as a full-length protein (it can be efficiently purified only as a complex with aPKC) (29). Many important cellular proteins are similarly challenging to purify due to obligate binding partners, unstructured regions, or a propensity to aggregate. Our success in quantitively studying the PAR-6/aPKC complex suggests that our approach will be applicable to a wide range of protein complexes that have been refractory to purification.

Our focus here has been to establish and validate the dynamic sc-SiMPull approach through studying control samples and the well-characterized aPKC/PAR-6 interaction. There are several features of this assay that we have not explored here but that will be valuable when working with complexes that are less stable and well-characterized. First, since it is now possible to begin counting protein complexes within seconds after lysis, this assay should be applicable to weak or transient protein-protein interactions that might be difficult to detect with other cell-based approaches. Second, by imaging and then immediately lysing cells or embryos, it will be possible to precisely (and rigorously) stage samples before lysis. This will allow investigators to measure how protein complexes are regulated across development or changes in cell state, revealing features of signal transduction that *in vitro* assays cannot capture. Finally, our framework should be readily generalizable to more than two fluorescent colors, allowing multi-protein assemblies to be studied with this approach. These ideas will be explored in future work.

## Supporting information

Key Resource Table

## Acknowledgments

We thank Luke Lavis for sharing JaneliaFluor dyes; Eric Drier and members of the Dickinson laboratory for helpful discussions and comments on the manuscript; and Robert J. Dickinson for assistance testing laser lysis equipment. This work was supported by a research grant from the Mallinckrodt foundation and by NIH R00 GM115964 and R01 GM138443 (DJD). DJD is a CPRIT Scholar supported by the Cancer Prevention and Research Institute of Texas (RR170054). Some strains were provided by the Caenorhabditis Genetics Center, which is funded by the NIH Office of Research Infrastructure Programs [P40 OD010440].

## Conflict of Interest

The authors declare that they have no conflicts of interests with the contents of this article.

## Author Contributions

DJD conceived of the project and built the laser lysis setup. SS and DJD together designed the experiments, performed the experiments, developed the analysis software, and analyzed the data. DJD wrote the first draft of the manuscript; both authors discussed and contributed to the final version.

## Materials and Methods

### Contact for Reagent and Resource Sharing

Requests for resources and further information should be directed and will be fulfilled by the Lead Contact, Daniel J. Dickinson.

### Experimental Model and Subject Details

*C. elegans* strains were fed *E*.*coli* OP50 and maintained on NGM growth medium at 20°C. The HaloTag-only control strain used in Figure S2 was constructed using Cas9-triggered homologous recombination with drug selection, as previously described (9).

### Method Details

#### Microfluidic device fabrication

The microfluidic device design used here was the same as in our previous work (3, 31). Device molds were made using SU-8 photolithography, following standard procedures. A 10:1 ratio of PDMS to curing agent (Sylgard 184 silicone elastomer kit, Dow Corning, Midland, MI) was mixed, poured onto the molds, and degassed for two minutes. The molds were then spin coated with the PDMS at 300 rpm for 30s. The PDMS was cured at 85°C for 20 minutes before peeling off the molds. A 2mm biopsy punch was used to create the inlet and outlet holes for each of the 12 channels per PDMS device.

24×60 mm glass coverslips were cleaned of visible dust using compressed nitrogen gas and placed in UV/Ozone cleaner for 20 minutes. Meanwhile, PDMS devices were prepared for glass bonding by air plasma treatment for 30s. Plasma treated PDMS devices were placed onto cleaned glass coverslips to create a permanent bond. Devices were immediately passivated by placing 2 µL of PEG solution (mPEG-Silane with 2% HPLC water and 0.02% Biotin-PEG-Silane) into the inlets. Once the PEG solution flowed to the ends of the channels, 0.5 µL of PEG solution was placed into the outlets. After a 30-60 minute incubation period at room temperature, the PEG solution was aspirated from the devices using a vacuum. The devices were rinsed twice by placing 2 µL of HPLC into both wells of each channel and vacuuming either the outlets or the inlets. The channels were dried thoroughly using a vacuum and placed into an airtight container containing desiccant. Devices were used within 1-2 months after PEG passivation.

Immediately prior to an experiment, devices were functionalized with anti-mNeonGreen nanobodies. The channels of the devices were rehydrated by placing 1.5 µL of SiMPull buffer (10 mM Tris pH 8.0, 50 mM NaCl, 0.1% Triton X-100, and 0.1 mg/mL BSA) into each well and vacuuming the inlets to remove most of the liquid while leaving buffer in the channels. A solution of 0.2 mg/mL Neutravidin in SiMPull buffer was prepared, and 1.5 µL of the solution was placed into each outlet. The Neutravidin solution was allowed to flow through the channel for a 10-minute incubation period at room temperature and washed with SiMPull buffer four times. A solution of 100 µM anti-mNeonGreen nanobody in SiMPull buffer was prepared, and 1.5 µL of the solution was placed into each outlet. The nanobody solution was allowed to flow through the channel for a 10-minute incubation period at room temperature and washed with SiMPull buffer four times. Any lane that was not immediately used was hydrated with 0.5 µL SiMPull buffer in each well and sealed with clear tape. When an embryo was ready to be placed into a channel, the tape was cut off using a scalpel. Occasionally, devices were functionalized and stored at 4°C overnight prior to an experiment.

#### Monovalent nanobody preparation

Monovalent nanobodies recognizing mNeonGreen were biotinylated via incubation with 100-fold molar excess of EZ-Link NHS-PEG_4_-biotin for 1 hour at room temperature. The reactions were quenched with Tris base and dialyzed to remove excess biotin. To determine concentration of final biotinylated mNeonGreen nanobody, UV absorbance at 280 nm was used.

#### HaloTag labeling

HaloTag ligands JF_646_, JFX_646_, and JFX_650_ yielded indistinguishable results and were used interchangeably in our experiments. Dyes were dissolved in acetonitrile and dispensed into 2 nmol aliquots. A speedvac was used to evaporate the solvent. The dried aliquots were stored at -20°C in dark, foil-wrapped containers. Immediately prior to labeling whole worms with HaloTag ligand, a dried aliquot was brought to a concentration of 1 mM with 2 µL DMSO. An E. coli OP50 culture was spun down and resuspended in ⅕ volume of S medium (150 mM NaCl, 1 g/L K2HPO4, 6 g/L KH2PO4, 5 μg/L cholesterol, 10 mM potassium citrate pH 6.0, 3 mM CaCl2, 3 mM MgCl2, 65 μM EDTA, 25 μM FeSO4, 10 μM MnCl2, 10 μM ZnSO4, 1 μM CuSO4). To obtain a final concentration of 10-15 μM HaloTag ligand, 1-1.5 μL of the DMSO aliquot was added to 100 µL of the bacterial suspension. Thirty microliters of the solution were placed into one well of a 96-well plate. About thirty worms at larval stage L4 were placed into the 30 μL of solution and incubated at 20°C overnight, shaking at 250 rpm. The following day, gravid adults were picked to be dissected in egg buffer (5 mM HEPES pH 7.4, 118 mM NaCl, 40 mM KCl, 3.4 mM MgCl2, 3.4 mM CaCl2).

#### sc-SiMPull from staged embryos

Gravid adults were dissected in egg buffer, and embryos were washed twice by sequentially transferring into two 40-100 µL drops of egg buffer using a mouth pipet. A zygote was then transferred to a 40-100 µL drop of regular or detergent-free SiMPull buffer, depending on the experimental conditions. Next, 0.5-1 µL of buffer was placed into the inlet well of a functionalized device. The zygote was transferred into the inlet well and either gently pushed into the channel using a clean 26G needle or gently pulled into the channel via vacuum. Channel flow was equilibrated to about 0.5 µL of buffer in each inlet by placing or removing buffer from the wells using a mouth pipette. Once the zygote was immobile and secure in the channel, two small pieces of clear tape were placed over the outlet and inlet wells, in that order, to prevent buffer evaporation and flow in the channel. The device, with the live zygote, was transferred to the TIRF microscope for data collection. Importantly, the embryo was not suffocated and able to continue developing up until the moment of laser lysis as PDMS is oxygen permeable and the non-oxygen-permeable tape was confined to covering the wells.

#### TIRF microscope construction

The custom-built microscope used for our experiments was built around an RM21 Advanced microscope platform (MadCity Labs, Madison, WI). The instrument is equipped with a piezo-Z stage for focusing; a 60X, 1.49NA Olympus APON TIRF objective lens; and micromirrors positioned below the objective for dichroic-free, multi-wavelength TIRF illumination (17, 18). Illumination light was provided by 488 nm and 638 nm LaserBoxx diode lasers (Oxxius, Lannion, France) which were focused through a 25 µm precision pinhole (Edmund Optics, Barrington, NJ) as a spatial filter, collimated, and focused again at the back aperture of the objective lens. The fluorescence signal, collected by the objective lens, was focused using a tube lens with a focal length of 250 mm, resulting in an initial magnification of 83.33X. The image was a split into green and far-red channels with a T565lpxr dichroic mirror (Chroma, Bellows Falls, VT), and the two images were collected side-by-side on the chip of a Prime95B scMOS camera (Teledyne Photometrics, Tucson, AZ). A set of relay lenses in the dual-view system introduced an additional 1.2X magnification, resulting in a total system magnification of 100X.

We made two modifications to the RM21 setup to enable dynamic sc-SiMPull experiments. First, we added a triggered 1064 nm pulsed laser (STP-40K, Teem photonics, Grenoble, France) which was installed on a separate optical path in order to focus at the sample plane rather than the back focal plane. The triggering pulses were provided by an Arduino Uno. The infrared laser was delivered to the objective via a short-pass dichroic mirror placed below the objective and the TIRF micromirrors (Figure 1B); this did not interfere with TIRF imaging, and allowed us to perform laser lysis and TIRF in rapid succession without any moving parts in the optical path. Second, we modified the MadCity Labs TIRF-lock system to enable continuous focusing even when the TIRF lasers were off. We installed a low-power (4.5 mW), 850 nm diode laser whose output was made co-linear with the TIRF lasers. After the beam exited the objective, the 850 nm light was separated from the TIRF excitation light via a dichroic mirror and focused onto a quadrant photodiode detector (MadCity Labs). The detector was linked directly to the piezo Z stage in order to maintain focus, via software provided by MadCity Labs. The entire hardware setup was controlled by Micro-Manager 2.0-gamma (https://micro-manager.org/).

#### Embryo lysis and TIRF microscopy

The channel in-use was centered on the objective and the embryo was located using transmitted light. Transmitted light was used to record a snapshot of the embryo for staging purposes. The stage was moved 120-200 µm away from the embryo to focus in TIRF. TIRF focus was determined using a 15 mW, 488 nm laser and a 50 mW, 638 nm laser, and TIRF lock was engaged. The TIRF lasers were turned off, and the stage was moved to center the embryo using transmitted light. The embryo was lysed using a 1064 nm Q-switched Nd:YAG pulsed laser. The stage was quickly moved 60-420 µm away from the point of lysis; the transmitted light was turned off, and continuous time lapse acquisition using TIRF excitation was performed for 10-15 minutes. A stopwatch was used to record the elapsed time between laser lysis and the start of TIRF acquisition. mNG was excited using 15 mW, 488 nm laser; HaloTag ligands JF_646_, JFX_646_, and JFX_650_ were excited using a 50 mW, 638 nm laser for all experiments except the lower laser power experiments for which 5 mW were used.

### Quantification and Statistical Analysis

#### Analysis of static SiMPull data

The analysis presented in Figure 2 was done exactly as in our previous work (3, 31) using our open-source SiMPull analysis software. Briefly, the first 50 frames of images from each stage position (along the length of the microfluidic channel) were averaged to produce high-contrast images for segmentation. Molecules in each frame were then detected using probabilistic segmentation (5) applied to both imaging channels. The left and right sides of the dual-view image were registered automatically using the built in `imregcorr` function in MATLAB. The software then checks for co-localization; signals that within one point spread function of each other were considered colocalized. The software automatically ignores images that have a density of spots greater than 0.3 per µm^2^, which we previously found is the maximum density below which <5% of spots colocalize by chance (3, 31). For the remaining images, colocalization was measured and summed across each sample by counting the number of colocalized molecules and dividing by the total number of molecules detected in each fluorescence channel. Data are presented as individual data points (one data point per embryo), with a weighted mean and 95% confidence interval calculated across the entire dataset. 95% confidence intervals were calculated using the formula

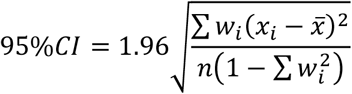

where *x*_*i*_ is the measured value from one single-embryo experiment, 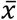 is the weighted mean, *w*_*i*_ is the weight (defined as the number of molecules counted in experiment *i* divided by the total number of molecules counted in all experiments), *n* is the number of experiments performed, and the sums are over all experiments.

#### Automated processing of dynamic SiMPull data

We added additional functionality for processing dynamic SiMPull data to our open-source SiMPull Analysis Software package, and then used these new tools to extract the relevant information about each observed molecule in our dynamic experiments. The algorithm is illustrated in Figure 3B. The user chooses the bait channel, which is the fluorescent protein that has been pulled down in the experiment (in the present work, this was always the green channel, since we exclusively used antibodies against mNG; however, this is flexible and other antibodies could be used if needed). The user also provides a reference image of either control proteins or beads, which is used to perform automated image registration of the left and right dual-view channels. Once registration is complete, the software bins each long TIRF movie into “windows” (typically 50 frames in length) that are averaged to produce high-contrast images for segmentation. Consecutive windows are subtracted to generate difference images, which are then subjected to probabilistic segmentation (5) to identify the locations of newly appearing signals. Segmentation is only done on the bait protein channel, since our goal is to detect appearing bait proteins and ask whether they co-appear with prey proteins. Once the locations of appearing bait molecules are identified, the software returns to the raw (non-binned) images and extracts the fluorescence intensity as a function of time, at each bait spot location, for both the bait and prey channels. Since our datasets can be as long at 16,000 image frames, analyzing the entire intensity trace for each molecule would be excessively costly (and is unnecessary). Instead, the software extracts and saves the 50 frames before a spot is expected to appear and the 450 frames afterwards (500 frames total). The intensity traces are analyzed using a Bayesian changepoint detection algorithm (23), with a log-odds ratio cutoff of 2 (corresponding to 99% confidence in each reported step) to find the time points when changes in intensity occur. The first increase in bait channel intensity is considered the time of spot appearance; traces where no change in intensity occurs within the first 100 frames are not analyzed further. Once the time of bait protein appearance is found, the prey channel is checked for a corresponding intensity increase at the same time. Prey spots that appear at the same location and within 4 frames (200 ms) of the bait protein are considered co-appearing. The software saves a data file that contains the location, time of appearance, and intensity trace for every detected molecule, along with whether it was called as co-appearing. This data file can then be queried further to extract the key information from each experiment.

#### Post-processing of dynamic SiMPull data

We created a set of tools within our software package to view the raw images, intensity-vs-time traces, co-appearance measurements and kinetic properties of our data. These scripts allow the user to inspect and evaluate the data quality, retrieve co-appearance information from the data file, apply filters for molecular density and blinking, and generate plots. All plots shown in this paper were generated using these open-source tools.

A typical workflow was to 1) inspect the images and ensure high-quality data; we excluded images with excessive background fluorescence (caused by poor coverslip cleaning) or for which no, or very few, fluorescent molecules appeared (this was sometimes caused by flow within the microfluidic channel due to poor sealing); 2) check images to ensure that the automated analysis delivered results that were reasonable; 3) correct image registration, if needed; and 4) perform downstream analyses of co-appearance vs. time, dwell time distributions, and density, depending on the experiment. We intentionally automated as much of the analysis pipeline as possible, in order to reduce user interventions that might introduce bias.

#### Kinetic analysis: Extracting *k*_*off*_ from co-appearance vs. time traces

To estimate protein complex dissociation kinetics from plots of co-appearance vs. time, we fit the raw data from each experiment to an exponential decay curve. We made two assumptions in our analysis. First, we assumed that PAR-6 and aPKC form a 1-to-1 heterodimer, which is supported by *in vitro* biochemical and crystallographic studies (24). Second, we assumed that the solution concentration of the bait protein (aPKC in our experiments) is small relative to the concentration of the prey protein. This assumption is common in receptor-ligand binding studies (30) and is justified in this particular case because the bait protein will be rapidly depleted from solution by binding to immobilized antibodies within the microfluidic device, while the prey protein remains free to participate in biochemical reactions in solution. With these assumptions, the relaxation of this protein-protein binding reaction to equilibrium will be described by the equation

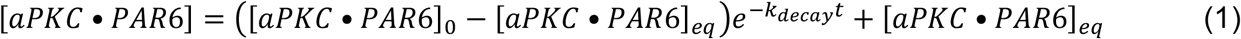

where *aPKC* • *PAR*6 denotes the aPKC/PAR-6 complex (we drop the hyphen from “PAR-6” for clarity), the ‘0’ subscript denotes the initial concentration, the ‘eq’ subscript denotes the concentration at equilibrium, and *k*_*decay*_ *= k*_*off*_ *+ k*_*on*_[PAR6] (30). Since aPKC was the bait protein in our experiments, what we measured was the fraction of aPKC molecules bound to PAR-6 – that is, the ratio of [PAR6/aPKC] to the total (bound + unbound) aPKC concentration [aPKC]_tot_. We denote the fraction bound as *F*. Dividing through by [aPKC]_tot_, equation 1 becomes

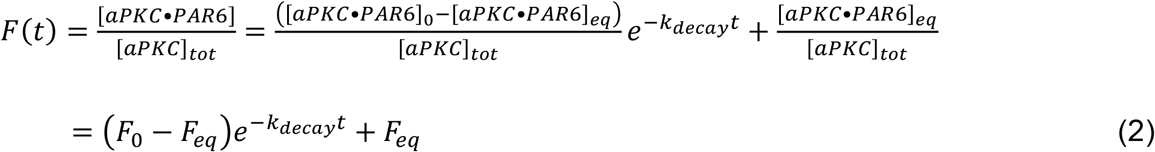

We fit each dataset to the equation *y = Ae*^*-Bt*^ *+ C* to extract the parameters *A = (F*_*0*_ *– F*_*eq*_*), B = k*_*decay*_ and *C = F*_*eq*_ using the built in `fit` function in MATLAB to perform a nonlinear least-squares fit. Fits with R^2^ < 0.2 were excluded from our analysis.

To estimate *k*_*off*_ for the complex, we derived an expression for *k*_*off*_ in terms of *k*_*decay*_ and *F*_*eq*_, which we obtained from our curve fits, as follows. Beginning with the expression for the equilibrium dissociation constant *K*_*D*_,

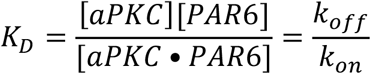

we rearrange to obtain

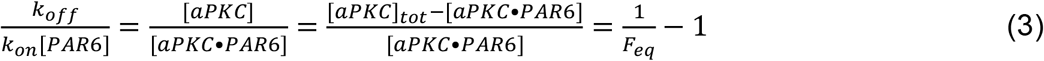

From the expression for *k*_*decay*_, we also have

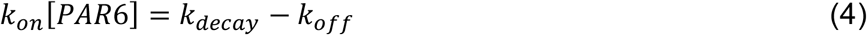

Substituting equation 4 into equation 3 and rearranging, we obtain our final expression for *k*_*off*_:

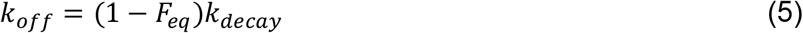

Equation 5 allows us to calculate *k*_*off*_ from our experimental co-appearance vs. time data. Note that under conditions where protein complexes that dissociate do not re-bind (for example, due to dilution), *F*_*eq*_ *=* 0 and equation 5 reduces to *k*_*off*_ *= k*_*decay*_, as expected. Our experimental fits yielded *F*_*eq*_ < 20% for all samples and *F*_*eq*_ < 5% for 10/12 samples, indicating that under our conditions, the contribution of rebinding to equilibration is minor and thus *k*_*off*_ will always be close to *k*_*decay*_.

One final complication is that our experimental estimates of *F*_*eq*_, and thus *k*_*off*_, will be affected by incomplete labeling of HaloTag with fluorescent dyes. Because *F*_*eq*_ was small in our experiments, the magnitude of the resulting error will also be small, but we corrected for it nonetheless. Based on an estimated labeling efficiency of ∼75% (Figure 3E), we divided *F*_*eq*_ by 0.75 before calculating *k*_*off*_.

#### Kinetic analysis: Extracting *k*_*disappear*_ from dwell time distribution analysis

To measure the dwell times of individual prey proteins, we wrote a simple post-processing script that calculates the interval between co-appearance of a complex and the time when the prey intensity returns to its initial (background) value. We deliberately neglected fluctuations in the prey protein intensity (as long as intensity remained above background), because we have observed that single HaloTag molecules labeled with JaneliaFlour dyes can fluctuate in intensity over time (DJD, unpublished observations). Our dwell time distribution analysis followed the template given in (28). We calculated a prey disappearance probability (per image frame) using the equation

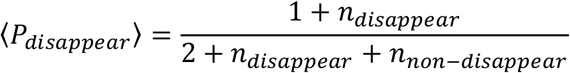

where *P*_*disappear*_ is the probability that the prey protein disappears (angular brackets denote the mean), *n*_*disappear*_ is the number of disappearance events observed, and *n*_*non-disappear*_ is the number of frames a prey molecule was observed to remain bound and not disappear. The disappearance probability is related to the disappearance rate constant by

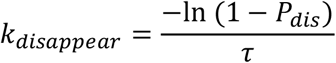

where *1τ* is the measurement interval (28) – typically 50 ms in our experiments. 95% confidence intervals around *k*_*disappear*_ were calculated from the beta distribution using the MATLAB function `betaincinv.` We report the confidence interval calculated in this way for each experiment, and a geometric mean and 95% confidence interval for *k*_*disappear*_ across each dataset. The geometric mean was chosen because normality tests indicated that our estimates of *k*_*disappear*_ were most likely to be log-normally distributed; normality testing and calculation of confidence intervals was done using Graphpad Prism software.

---

**Figure S1:**
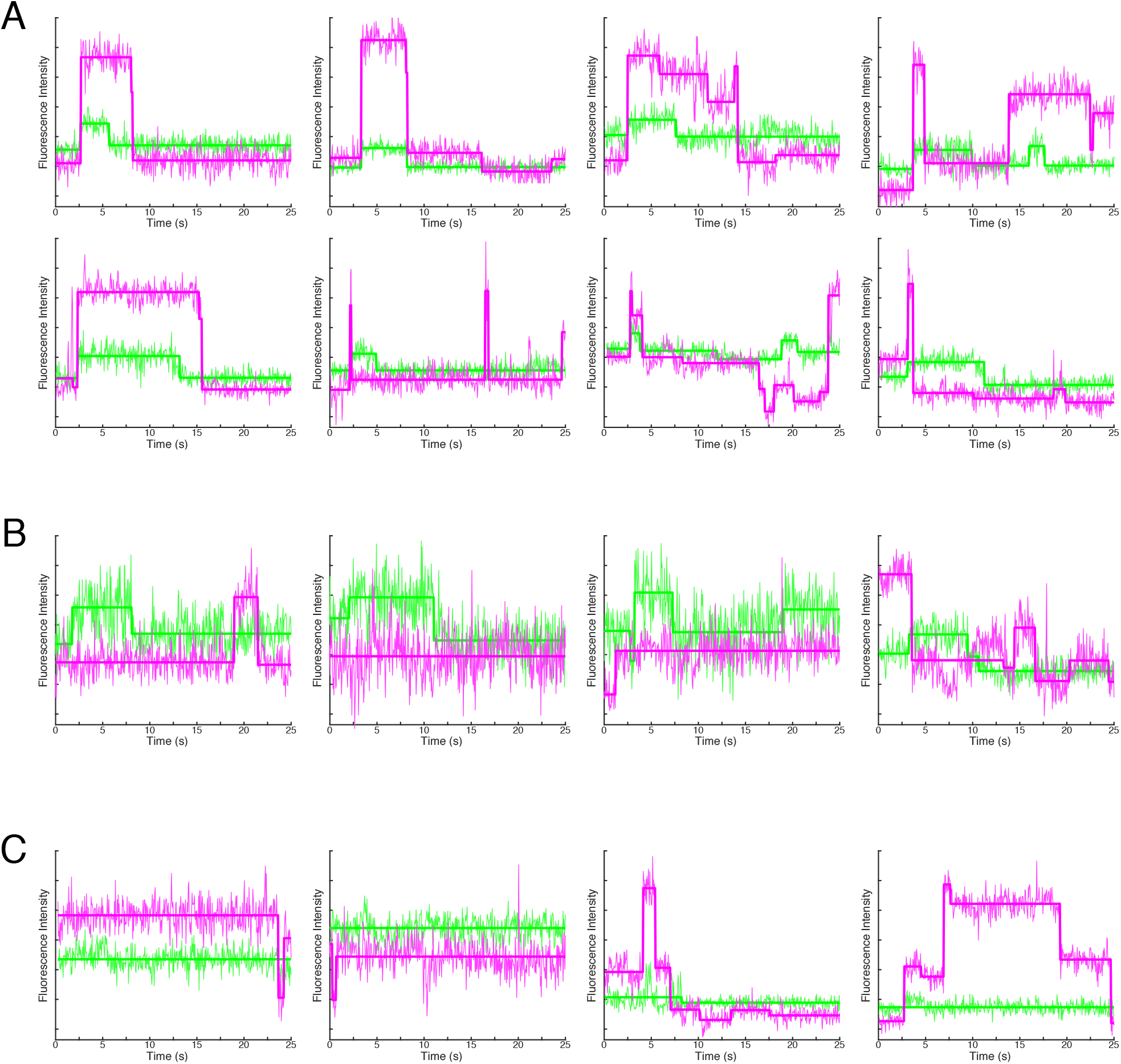
Example intensity vs. time traces for co-appearing and non-co-appearing molecules. A) Examples of traces that were classified as co-appearing. B) Examples of traces that were classified as not co-appearing. C) Examples of traces for which the bait protein could not be found. Most of these are likely mis-identified spots from the segmentation, and they were not considered in the analysis.

**Figure S2:**
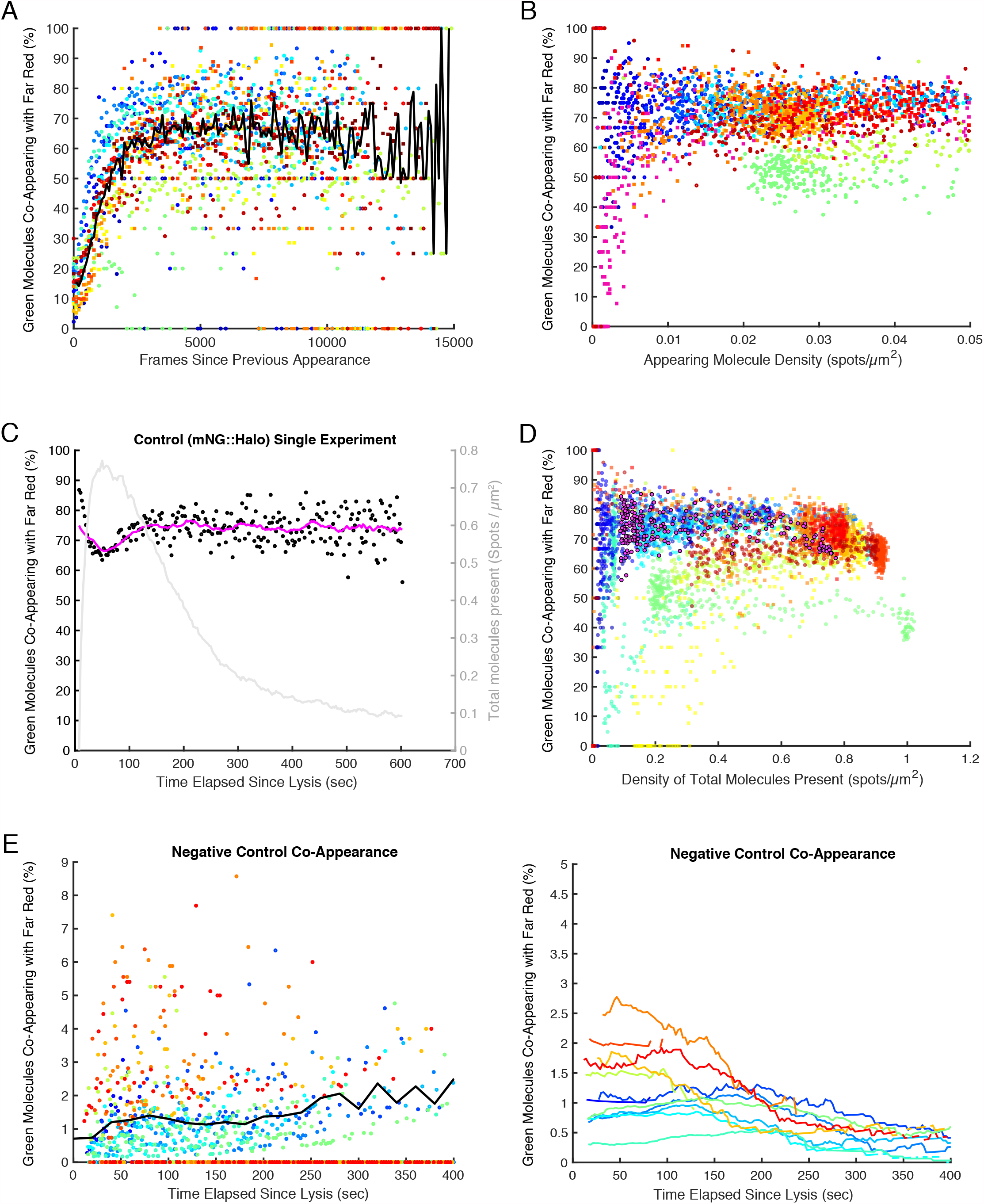
Sensitvity, dynamic range, specificity, and artifacts in dynamic sc-SiMPull. A) The effect of photo-blinking on detection of co-appearance. For each control mNG::HaloTag dataset (represented by a different color), data were filtered to identify spots that appeared at the same location as a previous spot. Then, co-appearance was calculated for just these spots, as a function of the time since their previous appearance. The sharp drop in co-appearance at short times indicates that many of these spots are blinking molecules that do not co-appear when they re-appear. Circles represent data that were acquired at 50 mW laser power, and squares represent data that were acquired at 5 mW laser power. N=14. B) The effect of low appearing molecule density on the precision of co-appearance measurements. Control mNG::HaloTag datasets were filtered to remove blinking spots, and then co-appearance was plotted as a function of the number of mNG molecules appearing in each 50-frame (2.5s) window. At very low densities – <0.01 molecules per µm^2^, which corresponds to only a few molecules per window – the measurements become noisy and exhibit horizontal “lines” (for example at 50%, 67% and 75% co-appearance) due to only a few molecules being counted in that window. Note that for clarity, the x-axis of this plot was truncated at 0.05 spots/µm^2^; windows that had a higher density of appearing molecules are not plotted. The complete datasets are shown in (D), below. Colors indicate different experiments. Circles represent data that were acquired at 50 mW laser power, and squares represent data that were acquired at 5 mW laser power. N=14. C) Co-appearance vs. time for a single control experiment that showed a pronounced dip in co-appearance at very high density. Black points show individual measurements of the fraction of molecules that co-appeared; each point corresponds to one 50-frame (2.5s) window of the data. The magenta curve is a moving average of these data. The gray curve shows the density of appearing spots as a function of time, plotted on the right Y axis. D) The effect of high molecule density, shown by plotting co-apperance as a function of number of molecules present in each average intensity projection. Colors indicate different experiments. Circles represent data that were acquired at 50 mW laser power, and squares represent data that were acquired at 5 mW laser power. The dataset shown in (C) is represented by magenta points outlined in black. N=14. E) Co-appearance vs. time for a set of experiments with a control, non-interacting protein pair (mNG::PAR-1 and HaloTag::GFP-nAb). Points on the left graph represent measurements of co-appearance for individual 50-frame (2.5s) windows, and the black trendline is a moving average across all datasets. The right graph shows moving averages for each dataset. Colors indicate different experiments. N=12.

## Supplemental Movie Legends

**Movie S1: Laser induced lysis of *C. elegans* zygote**

Brightfield view of a *C. elegans* zygote being lysed by a single shot from the pulsed infrared laser. The movie plays at actual speed.

**Movie S2: TIRF acquisition of the mNG::HaloTag fusion protein binding to the coverslip**

Time-lapse of mNG::HaloTag molecules appearing at the coverslip. Magenta is Halo ligand JFX_646_; Green is mNeonGreen. TIRF acquisition begins 11 seconds after cell lysis. The movie plays at 10 frames per second, and each frame is an average of 50 frames (1s) of actual data. Thus, each 1s of the movie represent 10s of actual data acquisition. Scale bar represents 10 µm.

**Movie S3: TIRF acquisition of mNG::aPKC and PAR-6::HaloTag molecules appearing at the coverslip**

Time-lapse of mNG::aPKC molecules binding to the coverslip and forming complexes with PAR-6::HaloTag molecules. Magenta is endogenous PAR-6::HaloTag bound to Halo ligand JFX_650_; Green is endogenous mNeonGreen::aPKC. TIRF acquisition begins 5 seconds after cell lysis. The movie plays at 10 frames per second, and each frame is an average of 50 frames (1s) of actual data. Thus, each 1s of the movie represent 10s of actual data acquisition. Scale bar represents 10 µm.

## Notes

### Competing Interest Statement

The authors have declared no competing interest.

